# Distinct Multivariate Structural Brain Profiles Are Related to Variations in Short- and Long-Delay Memory Consolidation Across Children and Young Adults

**DOI:** 10.1101/2021.08.24.457558

**Authors:** Iryna Schommartz, Philip F. Lembcke, Francesco Pupillo, Henriette Schuetz, Nina Wald de Chamorro, Martin Bauer, Angela M. Kaindl, Claudia Buss, Yee Lee Shing

## Abstract

From early to middle childhood, brain regions that underlie memory consolidation undergo profound maturational changes. However, there is little empirical investigation that directly relates age-related differences in brain structural measures to the memory consolidation processes. The present study examined system-level memory consolidations of intentionally studied object-location associations after one night of sleep (short delay) and after two weeks (long delay) in normally developing 5-to-7-year-old children (n = 50) and young adults (n = 39). Behavioural differences in memory consolidation were related to structural brain measures. Our results showed that children, in comparison to young adults, consolidate correctly learnt object-location associations less robustly over short and long delay. Moreover, using partial least squares correlation method, a unique multivariate profile comprised of specific neocortical (prefrontal, parietal, and occipital), cerebellar, and hippocampal subfield structures was found to be associated with variation in short-delay memory consolidation. A different multivariate profile comprised of a reduced set of brain structures, mainly consisting of neocortical (prefrontal, parietal, and occipital), and selective hippocampal subfield structures (CA1-2 and subiculum) was associated with variation in long-delay memory consolidation. Taken together, the results suggest that multivariate structural pattern of unique sets of brain regions are related to variations in short- and long-delay memory consolidation across children and young adults.

**RESEARCH HIGHLIGHTS:**

- Short- and long-delay memory consolidation is less robust in children than in young adults
- Short-delay brain profile comprised of hippocampal, cerebellar, and neocortical brain regions
- Long-delay brain profile comprised of neocortical and selected hippocampal brain regions.
- Brain profiles differ between children and young adults.

## 1 INTRODUCTION

### 1.1 Memory consolidation across development

Humans have an impressive capacity to store and retrieve memories of past experiences, consisting of unique temporal-spatial features, for years and even decades (Squire et al., 2015; Tulving, 2002). This is made possible due to memory consolidation, a dynamic and complex process through which acquired memory traces become long-lasting (Dudai, 2012; Moscovitch & Gilboa, 2021). From an ontogenetic perspective, the ability to retrieve long-term episodic memories emerges with the offset of childhood amnesia, i.e., the inability to recollect early life events, around four to seven years of age (Alberini & Travaglia, 2017; Bauer, 2007; Scarf et al., 2013; Tustin & Hayne, 2010). From there on, successful retrieval of complex memory representations starts to steadily improve (Drummey & Newcombe, 2002; Riggins, 2014; Sluzenski et al., 2006). In many nations, this age range is associated with the transition from kindergarten to school and is accompanied by dramatic increases in learning and knowledge accumulation, advancing cognitive functioning (Brod et al., 2017; McKay et al., 2022) and potentially also memory consolidation (cf. Nolden et al., 2021). However, little is known about the ability to consolidate memories over short and longer time in children who are about to start the school and face the necessity to retain plethora of newly acquired information.

Although much is known about how memory representations are encoded and retrieved in childhood, memory retention across longer consolidation periods (Murre & Dros, 2015; Ebbinghaus, 1885) is much less researched and may progress with different temporal dynamics in children who are about to start the school in comparison to adults (Peiffer et al., 2020; Wang et al., 2018; Wilhelm et al., 2008). For instance, it has been shown that short-delay memory consolidation rate (i.e., measured after one night of sleep) is comparable between children aged 6 to 8 years and young adults for word-pair associates (Wilhelm et al., 2008). On the other hand, it was also shown that in children and adolescents, successful retrieval of events over a longer time (e.g., one week) increases with increasing age (Østby et al., 2012). However, no study to date has directly compared memory consolidation of complex representations over short (i.e., one day) and long delays (i.e., weeks), and examined how maturational differences in brain structures between children and adults may account for potential age-related differences in memory consolidation. Therefore, in this study, we compared the retention rate of 5- to 7-year-old children and adults for learned object-location associations over one night as well as two weeks after encoding. Furthermore, we examined to what extent differences in retention rate are associated with multivariate patterns of structural measures of brain regions that are known to support memory consolidation.

### 1.2 Neural correlates of memory consolidation across development

Middle childhood is characterized by profound changes in cortical and subcortical brain regions related to mnemonic processes (Ghetti & Bunge, 2012; Ghetti & Fandakova, 2020; Lenroot & Giedd, 2006; Ofen, 2012; Ofen et al., 2007; Shing et al., 2010). For instance, the hippocampus, which is associated with the binding of event features into a coherent representation, reaches its relative maturity in late childhood/adolescence, depending on the subfields (Keresztes et al., 2017, 2022; Lee et al., 2014; Shing et al., 2008, 2010; Sluzenski et al., 2006). On the other hand, prefrontal brain regions, show protracted maturation into late adolescence/young adulthood (Gogtay et al., 2004; Muftuler et al., 2012; Sousa et al., 2018; Uda et al., 2015). This includes (i) the ventrolateral prefrontal cortex (vlPFC) and the orbitofrontal cortex (OFC) associated with strategy use that benefit memory formation and retrieval (Badre & Wagner, 2007; Kuhl et al., 2012; Østby et al., 2012), and (ii) the ventromedial prefrontal cortex (Brod & Shing, 2018; van Kesteren et al., 2012) and the rostral medial prefrontal cortex (Mella et al., 2021) that are important for schema-integration processes that benefit long-term consolidation. Similar, posterior parietal cortex (PPC), particularly its ventral part – precuneus, and lateral occipital cortex (LOC) (Nishimura et al., 2015; Simmonds et al., 2017) show more protracted development. PPC was shown to be involved in successful recollection of items with precise contextual details (DeMaster & Ghetti, 2013) and LOC was found to be associated with the reinstatement of object-related information upon retrieval (Grill-Spector et al., 2001; Karanian & Slotnick, 2015) and neural specificity of scene representation at retrieval in 8 to 15 years old children (Fandakova et al., 2019).

Beyond neocortical regions, entorhinal cortex (EC) being an input-output-hub for hippocampus-neocortical interactions plays a crucial role in memory trace strengthening (Reagh & Yassa, 2014; Takehara-Nishiuchi, 2014) and its structural maturity was related to memory performance (Daugherty et al., 2017; Keresztes et al., 2017). Parahippocampal gyrus (PHC) also supports spatial context-related associative recollection (Davachi et al., 2003; Milton et al., 2011; Ranganath & Ritchey, 2012) and was found to relate to subsequent memory recollection and long-term memory improvements in middle-late childhood (Ghetti et al., 2010). Finally, the cerebellum showed increased activation during retrieval of long-term episodic memories (Andreasen et al., 1999) and prefrontal-cerebellar circuits were also found to be related to declarative memory processes (Vecchi & Gatti, 2020), associative learning and recognition (Steinlin, 2007; Timmann et al., 2010).

In an exceptional study that examined consolidation and its brain correlates in participants aged 8 to 19 years, Østby et al. (2012) showed that a thinner OFC was associated with higher short-delay (30 minutes) recall, while larger hippocampal volumes were related to higher memory retention rates (1-week/30-min ratio) in a visuospatial task. These findings indicate that extended developmental trajectories of the neocortical regions and the hippocampus may affect the memory consolidation processes over short and long delay differentially in children, beyond their effects on encoding or retrieval. However, no other studies have directly compared short vs. long delay memory consolidation in children, particularly in the younger age range, and related these with structural brain measures.

### 1.3 The current study

In this study, we examined the consolidation of well-learnt object-location associations across short delay (after one night of sleep) and long delay (a 2-week-period) between learning and retrieval, comparing 5-to-7-year-old children to young adults, who served as a reference group with a mature memory consolidation system. We hypothesized no differences in short-delay memory consolidation between children and young adults (Peiffer et al., 2020; Wang et al., 2018), but less robust long-delay consolidation in children in comparison to young adults (Ghetti & Bunge, 2012; Lebel et al., 2012; Shing et al., 2010). Furthermore, we applied a partial least squares correlation analysis (PLSC) to map behavioural memory consolidation measures (i.e., retention rate, RR) onto multiple structural regions-of-interest (ROIs) reported previously to be involved in memory processes. This powerful statistical technique allowed us to overcome shortcomings of univariate approaches in light of highly correlated and interconnected brain ROIs and to identify brain profiles comprised of structural brain measures that are, in a multivariate pattern, associated with for short- and long-delay memory consolidation, respectively (Nestor et al., 2002). We hypothesized that a brain profile comprised of medial temporal lobe (MTL), cerebellar and neocortical (i.e. prefrontal, parietal, occipital) regions would be associated with variations in short-delay memory consolidation. This is because the availability of detail-rich representation of associative memories and the use of schema-integration and strategic control over memory should be beneficial. We also expected that a unique brain profile comprised of neocortical (i.e. prefrontal, parietal) and cerebellar brain regions would be associated with variations in long-delay memory consolidation, due to the importance of strategic control over memory traces with decaying perceptual representations.

## 2 METHODS

### 2.1 Participants

For the recruitment of children, 4000 general research invitation letters were sent to randomly selected families with 5-to-7-year-old children in Berlin, Germany, of whom 110 families expressed interest in participation. After screening, 63 typically developing children were recruited to participate in the study. 46 young adults were recruited to participate in the study through advertisement in the newspaper, on the university campus, and through word-of-mouth.

All participants had normal vision with or without correction, no history of psychological or neurological disorders or head trauma and were term-born (i.e., born after 37 weeks of pregnancy). We included only children and young adults with at least average IQ > 85. Thirteen children were excluded due to incomplete task execution and missing data (n=6) or technical issues during data acquisition (n=7). Seven young adult participants were excluded due to incomplete task execution and missing data (n=5), and identification as an extreme outlier (n=2) based on interquartile range for learning and consolidation behavioural measures (IQR; above Q3 + 3xIQR or below Q1 – 3xIQR (Hawkins, 1980)). The excluded participants were comparable in terms of age, sex, and socio-economic status to the final sample. In summary, the final sample size consisted of 50 typically developing children (20 female, mean age: 6.37 years, age range: 5.5 – 7 years), and 39 young adults (20 female, mean age: 25.44 years, age range: 21.3 – 30.8 years; Table 1). Structural T1-weighted brain images for volumetric assessments of cortical and subcortical brain regions as well as high-resolution structural hippocampal scans were acquired in all children and adults. After considering quality check (see details below) and technical errors, 46 children and 39 adults provided usable T1 images, and 46 children and 35 adults provided usable high-resolution hippocampal scan.

**Table 1.** Sample characteristics by age group (children, young adults) showing comparability between groups.

| Children<br>(CH; N = 50) |  | Young adults<br>(YA; N = 39) |  | Group effect<br>(CH vs YA) |  |
| --- | --- | --- | --- | --- | --- |
| *M* | *SD* | *M* | *SD* | *p-value* | *w*<sup>2</sup> |

| Demographic measures |  |  |  |  |  |  |
| --- | --- | --- | --- | --- | --- | --- |
| Age | 6.37 | .37 | 25.44 | 2.66 | *** | .97 |
| Sex (M/F) | 30/20 |  | 19/20 |  | - | - |
| IQ Score | 116.18 | 13.75 | 108.15 | 12.02 | *** | .08 |
| Socioeconomical Status |  |  |  |  |  |  |
| Income – Mother | 3.6 | 1.8 | - | - | - | - |
| ISCED – Mother | 6.1 | 1.5 | 4.1 | 1.8 | *** | - |
| Education Years - Mother | 19.7 | 4.2 | - | - | - | - |

*Notes.* Income is based on a 1-7 Scale (1 = less than 15.000€, 7 = more than 100.000€). CH = children; YA = young adults; ISCED = International Standard Classification of Education 2011 (1 = primary education; 8 = doctoral level; UNESCO Institute for Statistics, 2012); IQ = Intelligence Quotient based K-ABC (Kaufman & Kaufman, 2012) for children and WAIS-IV (Wechsler, 2012) for young adults; M = mean; SD = standard deviation; $w^2$ = omega squared; \* $p < .05$ ; \*\* $p < .01$ , \*\*\* $p < .001$ (significant difference); ns: non-significant difference.

All participants or their legal guardians gave written informed consent prior to participation. The study was approved by the ethics committee of the Goethe University Frankfurt am Main (approval E 145/18). The participants were compensated for participation in the experiment with 100 Euro.

### 2.2 Measures

#### 2.2.1 Object-location associations task

The stimuli for the object-location associations task were chosen based on the curriculum in social studies and science for the first and second grade of the German primary school (see similar procedure in Brod & Shing, 2019). Sixty different semantic themes (e.g., forest, farm, etc.) were chosen according to the ratings provided by four primary school teachers that assessed the familiarity of first graders with the topics. For each semantic theme, four scene pictures were combined with four thematically congruent object pictures, resulting in four unique object-location associations (see Fig. 1 for an example). We identified 18 possible areas in the scenes to place the objects, one of which was assigned to each object-location association (for more detailed information see Supplementary Methods section). We presented the task using Psychtoolbox-3 (Kleiner et al., 2007) software in Matlab 9.5, R2018b (MATLAB, 2018).

**Figure 1.**
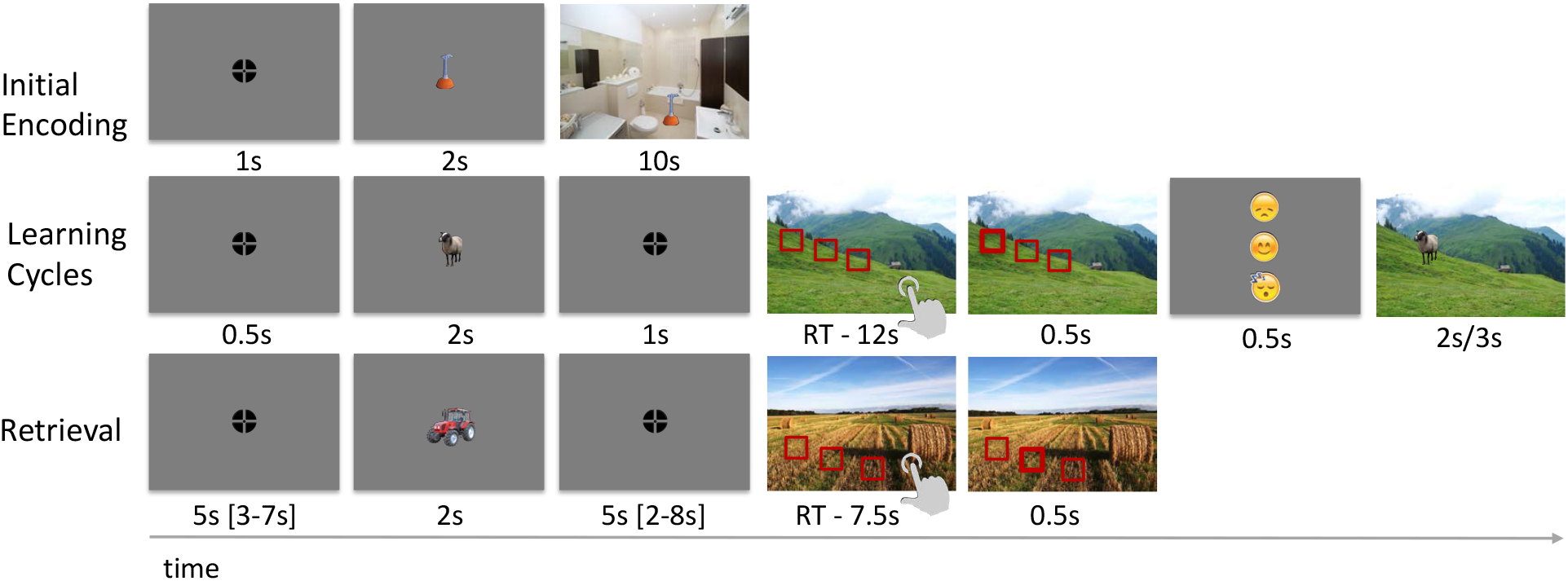
Experimental Task. (**A**) *Initial Encoding Phase*. Participants were instructed to remember 60 object-location pairs in total, memorizing the exact location of the object within the scene by creating a story or making a “mental photo” of the scene. (**B**) *Learning Phase*. Participants had to choose the correct location out of three choices and received feedback for their response. After the feedback, the correct object-location association was shown again. (**C**) *Retrieval Phase*. Participants had to choose the correct location of the object in the scene out of three options without feedback.

The task consisted of three phases as following (see Fig. 1):

i. Initial encoding phase (Day 0). For a set of 60 object-location pairs, participants saw the object followed by the same object superimposed on the scene at a particular location. Participants were instructed to remember 60 object-location pairs in total, memorizing the exact location of the object within the scene by elaboration (e.g., creating a story or making a “mental photo” of the scene), as such elaborative encoding strategies aid the recollection of the information (Craik & Tulving, 1975);
ii. Learning phase (Day 0). Participants learned the correct location of the object within the scene during adaptively repeated retrieval-encoding cycles (minimum two cycles, maximum four cycles). They had to choose the correct location out of three choices and received feedback for their response. After the feedback, the correct object-location associations were shown again. The cycles ended when participants provided correct responses to 83% of the trials or the maximum fourth cycle was reached.
iii. Retrieval phase (Day 1 and 14). Participants had to choose the correct location of the object in the scene out of three options without feedback. Note: The retrieval phase was carried out inside the MRI scanner with a functional sequence, of which the data is not included as we focused here on characterizing the retention rate behaviourally, both in terms of group comparison and relations to structural integrity.

#### 2.2.2 Assessment of demographic and cognitive covariates

In addition, IQ scores were assessed using the German version of the “Kaufman Assessment Battery for Children – Second Edition” (K-ABC II; Kaufman & Kaufman, N.L., 2015) in children and the “Wechsler Adult Intelligence Scale – Fourth Edition (WAIS-IV; Wechsler, 2015) in young adults. General socio-demographic questionnaires to assess socio-demographic characteristics of the participants were applied as well.

### 2.3 Experimental procedure

Testing took place across three days (see Fig. 2). On Day 0, the experimental procedure began with a short training to familiarize participants with the object-location associations task. Participants had to learn 60 object-location associations. The experimental task started with the initial encoding of the first 30 object-location associations. This was followed by a brief distraction task in which participants listened to and had to recall a string of numbers. This was followed by the learning phase with retrieval-encoding cycles until the 83% criterion was reached (or maximum of four cycles was reached). This procedure was done with the aim to minimize variances attributed to encoding, so that the comparison of subsequent memory consolidation could be made with starting points as similar as possible. After a short break, the same procedure was repeated with the other half of 30 object-location associations. On Day 1, short delay retrieval was conducted. Participants had to retrieve 30 object-scene associations learnt on Day 0. On Day 14, long delay retrieval was conducted. Participants had to retrieve another 30 object-scene associations learnt on Day 0.

**Figure 2.**
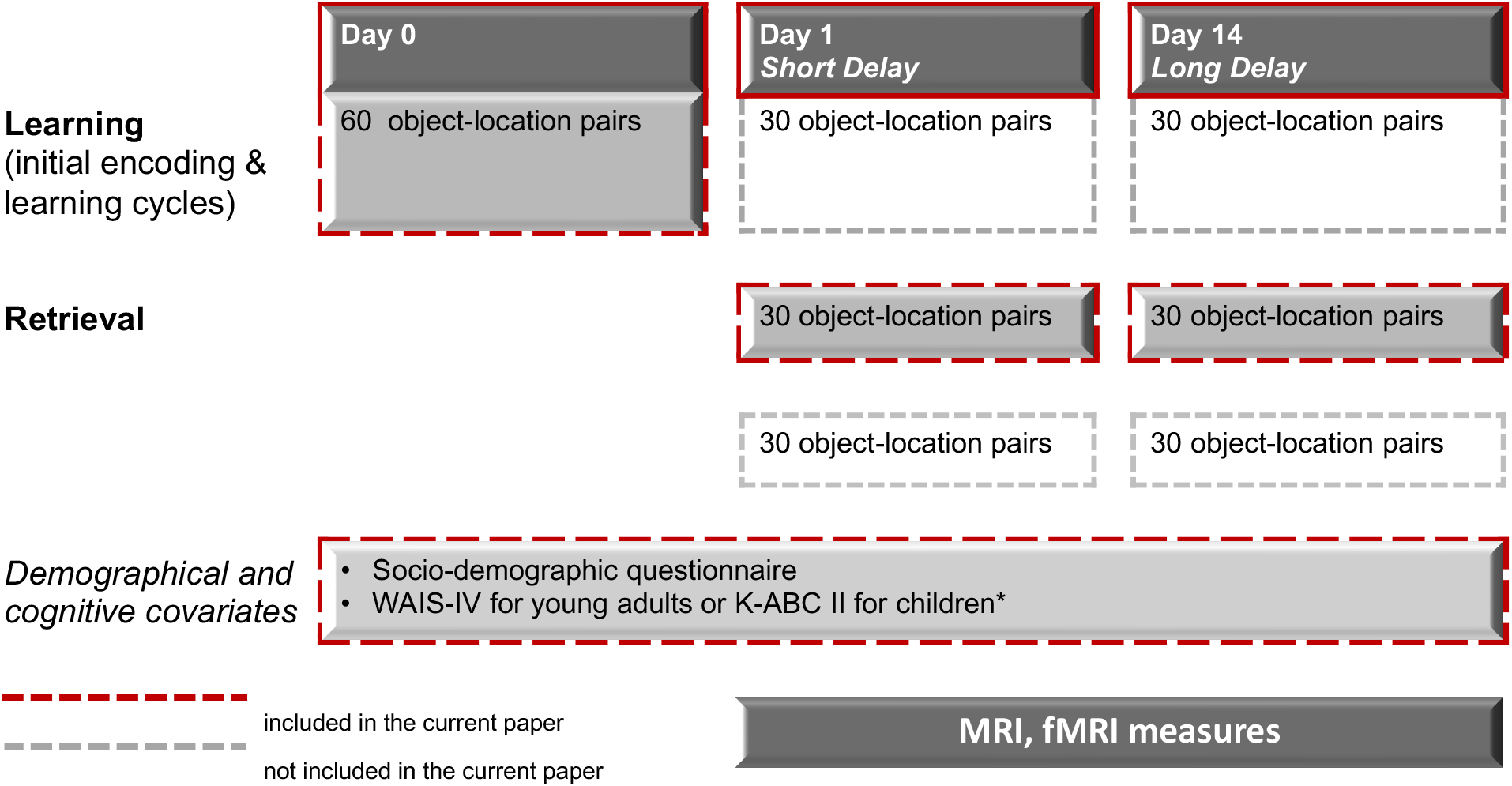
Experimental Procedure. The testing took place across three days. (i) On Day 0 participants had to learn 60 object-location associations. (ii) On Day 1 (short delay) the retrieval of 30 association pairs learnt on Day 0 was conducted. (iii) On Day 14 (long delay) the retrieval of another 30 association pairs learnt on Day 0 was conducted. Note that on Day 1 and 14, new sets of object-location pairs were learned to serve as baseline for fMRI analysis. Data on these newly learned pairs was not included here.

### 2.4 Magnetic resonance imaging

MR images were acquired on a 3-T SIEMENS PRISMA scanner (Siemens Medical Solutions, Erlangen, Germany) using a 64-channel head coil. An MPRAGE (magnetization prepared rapid gradient echo) T1-weighted sequence was applied with the following parameters: time repetition/time echo/time to inversion/Flip Angle = 2500 ms/2.9 ms/1070 ms/8°, matrix 256 × 256, field of view = 256. Each scan took 6 min 38 s. Each volume consisted of 176 sagittal slices with voxel sizes 1.0 × 1.0 × 1.0 mm.

### 2.5 Data Analysis

#### 2.5.1 Structural MRI data processing

##### Subcortical volumetric measures

Subcortical volumetric measures were derived using the anatomical pipeline of fMRIprep (version 20.2.1; Esteban et al., 2019), based on Nipype 1.5.1 (Gorgolewski et al., 2011). Here, brain tissue segmentation of cerebrospinal fluid, white-matter, and grey-matter was performed on the brain-extracted T1w scans using FAST (FSL 5.0.9; Zhang et al., 2001). Brain surfaces were reconstructed using recon-all (FreeSurfer 6.0.1; Dale et al., 1999). Volume-based spatial normalization to two standard spaces was performed through nonlinear registration with antsRegistration (ANTs 2.3.3), using brain-extracted versions of both T1w reference and the T1w template. Intracranial volume (ICV) was derived by the surfaced-based measures.

##### Cortical thickness measures

Cortical thickness measures were derived using the ABCD-HCP pipeline (Feczko et al., 2021) (https://github.com/DCAN-Labs/abcd-hcp-pipeline). The anatomical part of the pipeline includes three stages (refer to Supplementary Materials for more details). Cortical thickness measures were calculated from the distance between the reconstructed white matter and grey matter surfaces as well as from the reconstructed grey matter surface and cerebrospinal fluid boundaries. Structural data from one child participant was excluded due to poor quality assessed with the Qoala-T tool (Klapwijk et al., 2019).

##### Segmentation of hippocampal subfields

To delineate regions within the hippocampus, the Automated Segmentation of Hippocampal Subfields (ASHS) software (Yushkevich et al., 2015) was used, with a lifespan atlas created from manual segmentations (Bender et al., 2018).The approach used has been shown to be reliable and valid for delineating hippocampal subfields in a pediatric sample, including 6-14-year-old children (Bender et al., 2018). ASHS uses a multi-atlas method, integrating data extracted from segmentations of several hippocampi. Three regions of the hippocampal subfields within the hippocampal body were identified: the subiculum, Cornu ammoni regions 1 and 2 (CA1-2), and a region including CA3 and the dentate gyrus (CA3-DG). The CA3 and DG were not separated because even with high-resolution images the validity of their separation is not confirmed yet (Wisse et al., 2017). Presubiculum, subiculum, and parasubiculum were aggregated into the “Subiculum” subfield, and CA1 and CA2 were also collapsed into one single region for the same reason. The segmentation of the subfields was carried out only in the body of the hippocampus because the validity of the segmentation in the head and the tail is questionable (Wisse et al., 2017). The entorhinal cortex was also delineated on 6 consecutive slices anterior to the hippocampal body (Keresztes et al., 2020). Ranging of the hippocampal body was carried out following recent progress made towards the development of a standardized procedure for finding valid landmarks (Olsen et al., 2019).

##### Structural ROIs

The selection of structural ROIs was based on the review of the developmental literature on memory consolidation and related retrieval processes including structural MRI studies (Østby et al., 2012), which identified the involvement of OFC and hippocampus, as well as functional MRI studies (Andreasen et al., 1999; Davachi et al., 2003; DeMaster & Ghetti, 2013; Fandakova et al., 2019; Grill-Spector et al., 2001; Karanian & Slotnick, 2015; Kuhl et al., 2012; Mella et al., 2021; Milton et al., 2011; Nishimura et al., 2015; Ranganath & Ritchey, 2012; Reagh & Yassa, 2014; Simmonds et al., 2017; Steinlin, 2007; Timmann et al., 2010; van Kesteren et al., 2012), which identified the involvement of vmPFC, vlPFC, rostral medial PFC, precuneus, LOC, EC, cerebellum, PHG in memory retrieval. Based on the findings that hippocampal subfields may follow different developmental trajectories (Keresztes et al., 2017, 2022) and be differentially involved in memory delays (Atucha et al., 2021), EC and hippocampal body subfield volumes were also included as separate ROIs. The following corresponding regions of interest (ROI) were identified according to Desikan-Killiany atlas (Desikan et al., 2006) for the (i) volumetric output: cerebellar cortex, EC and hippocampal body subfields volumes (i.e., DG-CA3, Subiculum (Sub), and CA1-2); (ii) cortical thickness output: inferior frontal cortex (IFG; comprised of pars opercularis, pars triangularis and pars orbitalis), medial OFC, lateral OFC, rostral middle frontal cortex, praecuneus, superior parietal cortex, inferior parietal cortex, parahippocampal cortex, and lateral occipital cortex. As we did not have specific hypotheses based on prior research related to lateralization, these ROIs were collapsed across hemispheres for all following analyses (Dick et al., 2022). To control for differences in head size, brain volumes were adjusted for intracranial volume (ICV) using a covariance approach (Clifford et al., 1989; Raz et al., 2005; Voevodskaya et al., 2014). Cortical thickness was not adjusted for head size because cortical thickness and head size are not associated (Barnes et al., 2010; Mills et al., 2016).

#### 2.5.2 Behavioural Data Analysis

The analyses of all behavioural measures were performed with R packages (Version 4.0.4, R Core Team, 2021) in R Studio 1.4.1106 (RStudio, Inc.). Throughout the analyses, p-value significance levels were set to a < .05. We conducted a linear mixed-effect model for memory measures (accuracy defined as percentage of correct responses) using the lmer function from the lme4 package in R (Bates et al., 2015) and lmerTest (Kuznetsova et al., 2017). The linear mixed effect model was calculated with maximum-likelihood estimation and *Subject* as random intercept to account for between-subject variability in memory accuracy. As fixed factors, we included the within-subject factor of *Session* (short delay and long delay relative on correctly recalled items on Day 0) and the between-subject factor of *Group* (children and young adults). In addition, *IQ, Sex*, and *Handedness* (Kang et al., 2017; Willems et al., 2014) were added as covariates into the model. The main effects were followed up with Sidak post-hoc multiple comparisons. For group differences in memory measures, we conducted one-way independent analysis of variance (ANOVA). In case of violated assumptions of homogeneity of variances, a Games-Howell test was performed (Lee & Lee, 2018). The effect size estimation was performed with omega squared (w^2^) as a less biased estimate for reporting practical significance of observed effects (Finch & French, 2012; Okada, 2013; Troncoso Skidmore & Thompson, 2013). To determine the amount of variance explained by the model, we used partR2 package in R (Stoffel et al., 2020) with bootstrapping to calculate confidence intervals.

#### 2.5.3 PLSC (Partial Least Square Correlation): Linking Brain Structures to Behavioural Measures

We applied a multivariate Partial Least Square Correlation (PLSC) method (Abdi & Williams, 2013; Keresztes et al., 2017; Krishnan et al., 2011; McIntosh et al., 1996) to investigate the multivariate associations between predefined ROIs and variations in short- and long-delay memory retention rate (RR), both within and across both age groups. We extracted a latent brain profile that maximally shares common variance with either short-delay or long-delay variations in memory RR, in which a large part of the variance is driven by age-related differences in memory RR. We postulated that this multivariate approach is better suited than univariate assessment of the relation of different ROIs to memory consolidation due to their interconnectedness and interdependence (see e.g.,Genon et al., 2022).

First, we calculated a between subject correlation matrix between (i) a n x 16 matrix of volumetric or cortical thickness measures of all ROIs and (ii) an n-vector representing a continuous measure of either short-delay or long-delay RR: R = CORR (RR, ROIs). All measures entering the correlation were normalized. Singular value decomposition was used to decompose this correlation: R = USV’ into singular vectors U and V or saliences (Abdi & Williams, 2013; Krishnan et al., 2011). The left singular vector represents the short- or long-delay RR weights (U), the right singular vector of ROI weights (V) that represents the characteristic of brain structures that best represent R, and S is a diagonal matrix of singular values.

After that PLSC searches for a single estimable latent variable (LV) finding pairs of latent vectors with maximal covariance in a least-squares sense that represent the association between RR and ROI characteristics. Hence, LV represents distinct profiles of ROIs that have the strongest relation to either short- or long-delay RR. In addition, we calculated for each subject a single value, so-called within-person “short-delay brain score” and “long-delay brain score”, which are summary units of within-person robust expression of the defined LV’s profile. For this purpose, the model-derived vector of ROI weights (V) was multiplied by within-person vector of estimates of ROI structural measures.

We ran 5000 permutation tests to obtain a p-value to identify the generalizable vector of saliences or a LV and to assess whether the identified association is significant. Further, we identified the stability of within-LV weights through the subsequent bootstrapping on 5000 bootstrap resamples of the data and obtained a bootstrap ratio (BSRs) by dividing each ROI’s salience by its bootstrapped standard error. The BSRs are akin to Z-scores and considered to be normalized estimates of robustness (Keresztes et al., 2017), therefore when values are larger/smaller than ±1.96 (a < .05) their corresponding saliences are treated as significantly stable. Due to a single analytic step in multivariate statistical assessment using PLSC, no correction for multiple comparisons across all ROIs is necessary (McIntosh & Lobaugh, 2004).

Considering that more robust brain-wide association are observed in multivariate vs univariate methods (Marek et al., 2022), several merit can be highlighted for the application of PLSC to identify the relationship between specific MRI structural measures of brain anatomy and memory consolidation measures across age groups. The measures of brain anatomy are highly correlated and interconnected, specifically HC subfields, which may otherwise lead to statistical multicollinearity and redundancy, potentially reducing the statistical power to reveal neural-behavioural relationships when applying canonical statistical methods. Addressing these shortcomings, PLSC provides a statistically powerful technique which allowed us to map memory consolidation scores onto predefined multiple structural MRI ROIs (Nestor et al., 2002). It is important to note that our approach targets how brain structures are related to variations (or individual differences) in memory consolidations, and not of their involvement in consolidation processes in a within-person sense.

## 3 RESULTS

### 3.1 Behavioural results

#### 3.1.1 Learning Accuracy on Day 0

In the following we first tested for group-related differences on final learning accuracy of object-location associations during the learning phase on Day 0. To reach at least the set criterion of 83% correct responses, children needed significantly more learning-retrieval cycles on average (mean = 2.58, SD = .70, range: 2-4), in comparison to young adults who needed only 2 cycles as revealed by the Games-Howell test, b = −.58, p < .0001, 95% CI [−.77 – −.38]. Next, final learning accuracy defined as percentage of correctly retrieved locations of the objects within the scenes was separately calculated for items to be retrieved on day 1 and day 14 as they differed between participants (see Fig. 3A). The Games-Howell tests revealed no difference in final memory accuracy (i) between short delay (mean = 90.40; SD = 6.51) and long delay items (mean = 90.13; SD = 5.51) in children (see Table 2), b = −.003, p = .823, 95% CI [−.03 – .02]; ii) and between short delay (mean = 97.18; SD = 3.55) and long delay items (mean = 98.46; SD = 2.74) in young adults, b = .013, p = .078, 95% CI [−.001 – .02]. In addition, the Games-Howell test revealed a significantly higher percentage of correctly retrieved locations of the objects within the scenes in young adults in comparison to children, b = .076, p < .0001, 95% CI [.06 – .09]. Hence, despite our training-to-criterion procedure, young adults showed better memory performance than children at the end of the training. Observed groups differences in the final learning performance were considered in the subsequent modelling approach, which concentrated only on the items that were correctly retrieved during final learning performance.

**Figure 3.**
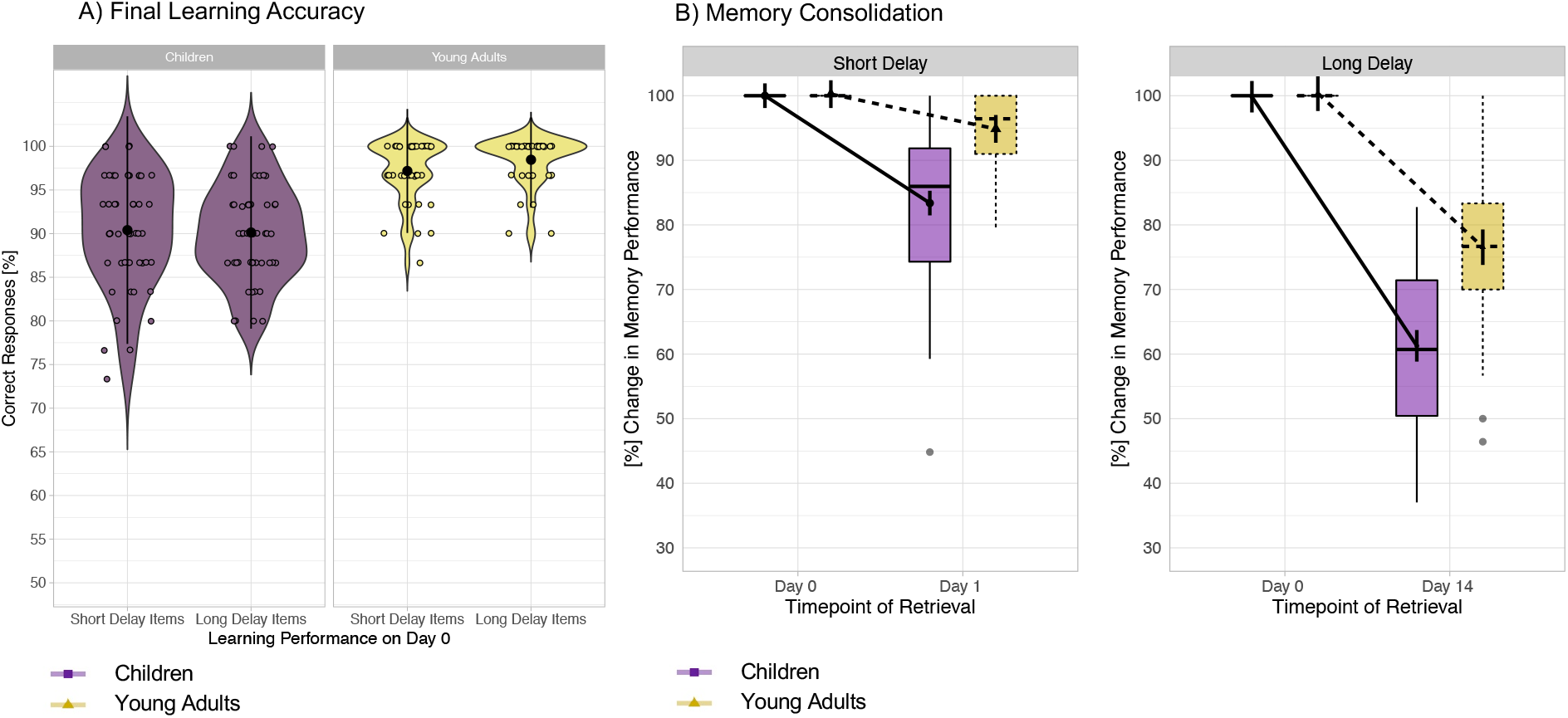
Final learning accuracy, and memory consolidation. (**A**) Final learning accuracy shows the percentage of correct responses for short delay items and long delay items after learning was completed in children and young adults. There was no difference in final learning accuracy for short and long delay item in either age group. Children needed between two to four learning-retrieval cycles to reach the criterion of 83% correct responses; while young adults need on average two cycles; (**B**) Memory consolidation over the course of two weeks, operationalized as percentage of correctly retrieved object-location associations on day 1 for items that were correctly retrieved on day 0 (after one night of sleep) for short delay, and percentage of correctly retrieved object-location associations on day 14 that were correctly retrieved on day 0 (after two weeks) for long delay. Error bars indicate model-based standard error. \**p* < .05. \*\**p* < .01. \*\*\**p* < .001(significant difference), ns: non-significant difference. P-values use Sidak correction for multiple comparisons.

**Table 2.**
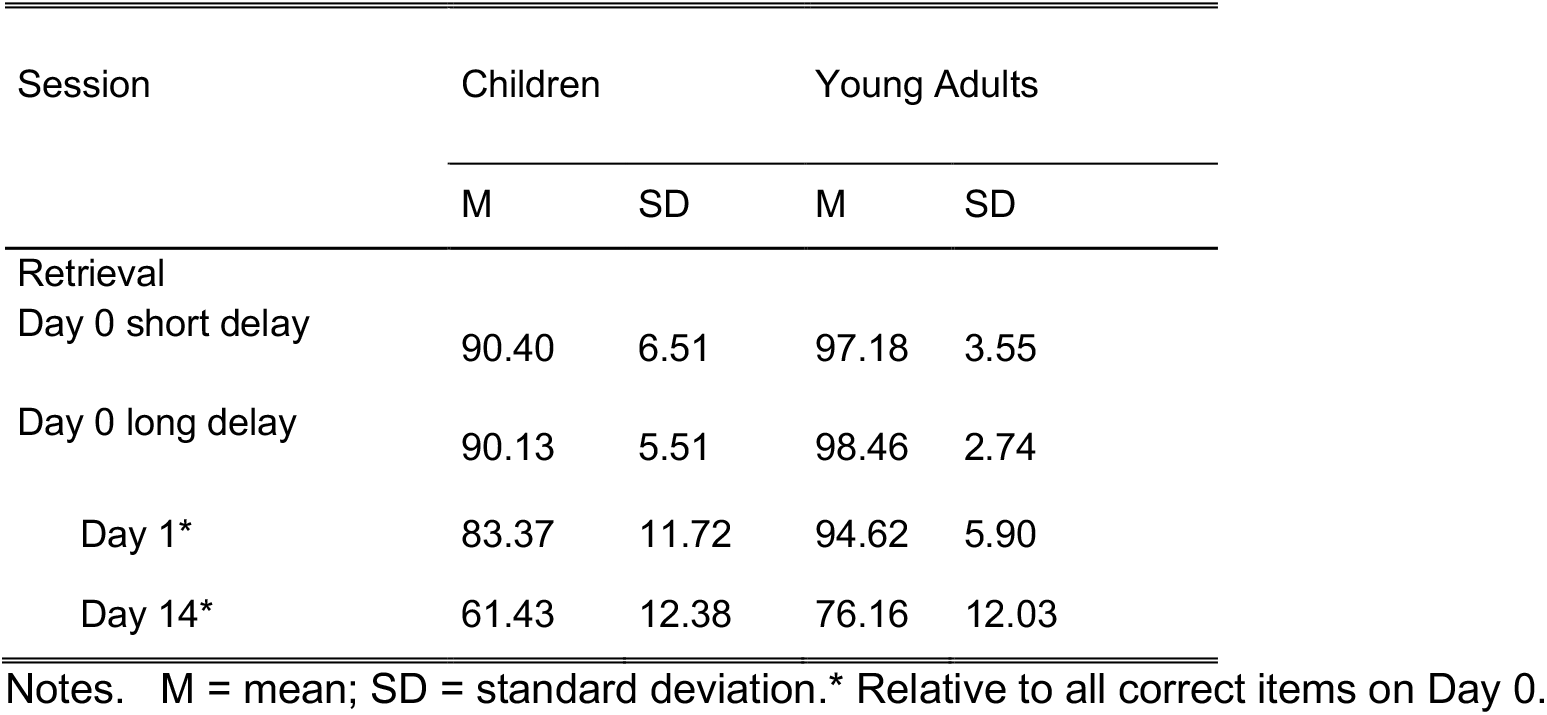
Descriptive statistics of memory performance based on percentage of correct answers by age groups.

#### 3.1.2 Memory Consolidation

In the following, we examined change in memory performance for correctly remembered items on day 0 across time, particularly testing for group differences in short- and long-delay memory consolidation (see Fig. 3B). The linear mixed-effects model for retrieval accuracies of learned object-location pairs explained a significant amount of variance, R^2^ = .74, 95% CI [.70 – .79]. We observed a significant main effects of *Session*, F_(2,178)_ = 342.05, p <.0001, w^2^ = .79, *Group*, F_(1,89)_ = 51.46, p <.0001, w^2^ = .36, and *Session* x *Group* interaction, F_(2,178)_ = 20.24, p < .0001, w^2^ = .18 (Supplementary Table S2 for a full overview). To interpret the interaction, we examined short delay and long delay separately (see Fig. 3C). For short delay, the model revealed a significantly steeper slope of accuracy decline from day 0 to day 1 in children in comparison to young adults (b = 11.25, t_(182)_ = 4.60, 95% CI [4.5 – 18.0], p < .0001), indicating less robust short-delay memory consolidation in children compared to young adults. Model-based Sidak post hoc comparisons for short delay revealed that (i) retrieval accuracy after one night of sleep declined significantly in children (b = 16.63, t_(182)_ = 10.26, 95% CI [12.2 – 21.1], p < .0001), and in adults (b = 5.38, t_(182)_ = 2.93, 95% CI [.3 – 10.4], p = .029). For long delay, the model also revealed significantly steeper slope of accuracy decline from day 0 to day 14 in children in comparison to young adults (b = 14.73, t_(182)_ = 6.02, 95% CI [7.9 – 21], p < .0001), indicating less robust long-delay memory consolidation in children. Sidak post hoc tests revealed a significant decline in long-delay retention rates in both groups (all p < .0001; Fig. 3B). Taken together both age groups showed a decline in memory performance over time, however, compared to young adults, children showed a steeper slope of memory decline for both short and long delay.

### 3.2 Brain-behavioural relationships

#### 3.2.1 Unique multivariate brains profiles are associated with short and long delay memory consolidation

As the next step, we applied PLSC to identify unique brain profiles of structural brain measures in relation to either short- or long-delay memory consolidation for the items that were correctly learnt on day 0, estimating the brain-behaviour pairings that covary together within and across age groups. The cross-correlational matrix was reduced to a set of single latent variables (LV) or saliences. Within age groups, we could not identify a single reliable LV to reliably represent brain-behaviour association (all p ≥ .43), due to the narrow age range within the groups and little age-related structural variability in all ROIs within the groups (Supplementary Table S3 and Figure S5 for a full overview). In the following the results across age groups will be presented. First, for short delay, we obtained a single composite score that captures individual differences in brain structures across both age groups in relation to memory RR, referred to in the following as “short-delay brain score”. With permutation test of significance, we identified a single reliable LV (p < .0001) that optimally represents an association between a profile of ROIs and short delay RR (r = .41). Using BSR that expresses the consistency with which the salience of a particular ROI is non-zero across subjects, we identified several stable components within the multivariate profile (see Fig. 4A): a positive short-delay RR association with ROI volumes of the cerebellum (BSR = 2.76, r = .26), ERC (BSR = 2.36, r = .27), and all hippocampal subfields: Sub (BSR = 2.82, r = .28), DG-CA3 (BSR = 2.73, r = .25), CA1-2 (BSR = 2.98 r = .28); a negative short-delay RR association with cortical thickness measures in parsopercularis part of the IFG (BSR = −5.85, r = −.42), parsorbitalis part of the IFG (BSR = −3.90, r = −.33), parstriangularis part of the IFG (BSR = −3.13, r = −.31), the lateral (BSR = −4.76, r = −.38), the medial (BSR = −5.89, r = −.43) the orbitofrontal cortex, the rostromedial cortex (BSR = −5.45, r = −.40), the precuneus (BSR = −4.00, r = −.35), the superior parietal cortex (BSR = −5.06, r = −.36), the inferior parietal cortex (BSR = −4.71, r = −.35), and the lateral occipital cortex (BSR = −3.26, r = −.32). Taken together, these stable components of the LV express the amount of information common to short-delay RR across both age groups and multivariate pattern of ROIs in specific neocortical, cerebellar, and hippocampal subfields structures (see Fig. 4B).

**Figure 4.**
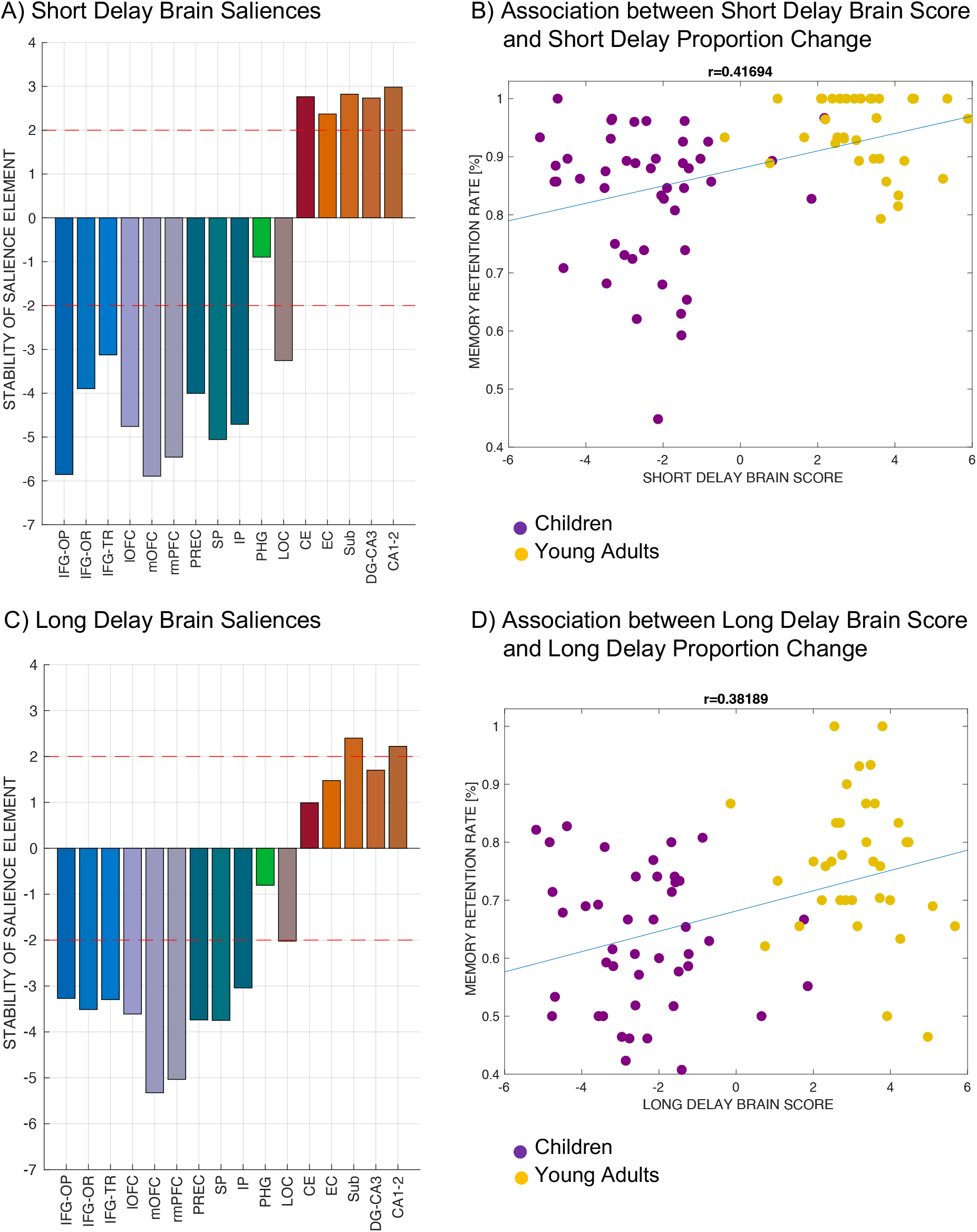
Multivariate profiles of brain structures associated with variations in memory consolidation. A) *Short Delay Brain Saliences*. Brain saliences or latent variables weights for each ROI were incorporated in the analysis to create one latent variable that expresses a composite short-delay brain score. Stability of salience elements is defined by Z-scores (depicted as red line: a value larger/smaller than ±1.96 is treated as reliably robust at (a < .05). B) *Association between Short Delay Brain Score and Short Delay Memory Retention Rate*. Brain short delay score or a latent variable plotted against short delay retention rate. There were significant age-related differences in short delay brain score, paralleling age-related differences in short delay retention rate. C) *Long Delay Brain Saliences*. Brain saliences or latent variables weights for each ROI were incorporated in the analysis to create one latent variable that expresses a composite long-delay brain score. Stability of salience elements is defined by Z-scores (depicted as red line: a value larger/smaller than ±1.96 is treated as reliably robust at (a < .05). D) *Association between Long Delay Brain Score and Long Delay Retention Rate*. Brain long delay score or a latent variable plotted against short delay retention rate. There were significant age-related differences in long delay brain score, paralleling age-related differences in short delay retention rate. *Note:* IFG – inferior frontal gyrus; OP – parsopercularis; OR – parsorbitalis; TR – parstriangularis; lOFC – lateral orbitofrontal cortex; mOFC – medial orbitofrontal cortex; rmPFC – rostromedial cortex; PREC – precuneus; SP – superior parietal cortex; IP – inferior parietal cortex; PHG – parahippocampal cortex; LOC – lateral occipital cortex; CE – cerebellum; EC – entorhinal cortex; SUB – Subiculum; DG-CA3 – dental gyrus and CA3; CA1-2 – CA1 and CA2 subfields of hippocampus. For group differences for each structural ROI (Houston et al., 2013), please refer to Table S8 in Supplementary Materials.

Second, with long delay, we obtained a single composite score that captures individual differences in brain structures across both age groups that relates to memory RR, referred to in the following as “long delay brain score”. We identified a single reliable LV (p = .0002) that optimally represents an association between predefined ROIs and long-delay RR (r = .382). BSR identified several stable components within the multivariate profile (see Fig. 4D): a positive long-delay RR associations with ROI volumes of hippocampal subfields: Sub (BSR = 2.40, r = .28), CA1-2 (BSR = 2.22 r = .28); a negative long-delay RR association with cortical thickness measures in the parsopercularis part of the IFG (BSR = −3.27, r = .23), the parsorbitalis part of the IFG (BSR = −3.51, r = .23), the parstriangularis part of the IFG (BSR = −3.30, r = .23), the lateral (BSR = −3.61, r = .29), the medial (BSR = −5.32, r = .40) the orbitofrontal cortex, the rostromedial cortex (BSR = −5.03, r = .38), the precuneus (BSR = −3.74, r = .28), the superior parietal cortex (BSR = −3.75, r = .26), the inferior parietal cortex (BSR = −3.04, r = .31), and the lateral occipital cortex (BSR = −2.22, r = −.32). These stable components of the LV express the amount of information common to long-delay RR across both age groups and multivariate pattern in neocortical and hippocampal ROIs. In contrast to short delay, cerebellar volumes and ECR and DG-CA3 hippocampal volumes do not contribute reliably to long-delay RR. The identified LVs account for a moderate portion of brain-behaviour covariance (short delay: 41%, long delay: 38%). Of note is that not all included ROIs contributed significantly to the covariance with memory performance, indicating specificity within profiles of brain ROIs with relation to either short- or long-delay memory RR (see Fig. 4D).

In addition, when testing for age differences in the brain scores, t-test revealed that the short-delay brain score, t_(78,68)_ = −17.17, p < .0001, as well as the long delay brain score, t_(78,95)_ = −18, p ≤ .0001, were significantly higher in young adults in comparison to children. This suggests that while the brain profiles were identified pulling across both age groups, there are age differences in the derived brain score, in parallel to age differences in behavioural memory consolidation measures.

## 4 DISCUSSION

In the present study, we investigated memory consolidation of correctly learned object-location associations (through adaptive and intentional learning) after short delay (with one night of sleep) and long delay (after 2 weeks), comparing performance in 5-to-7-year-old children and young adults. We found that: (i) children, in comparison to young adults, showed less robust memory consolidation of correctly learnt associations both after one night of sleep and after a 2-week-period; (ii) applying multivariate PLSC analysis with structural high-resolution MRI data, we identified a) a stable brain profile comprised of neocortical (prefrontal, parietal, and occipital), cerebellar, and MTL (i.e., all hippocampal subfields and EC) structures that is associated with variations in short-delay memory consolidation across both age groups; b) a stable brain profile comprised of more specific neocortical (prefrontal, parietal and occipital), and MTL (i.e., subiculum and CA1-2 hippocampal subfields) structures that is associated with variations in long-delay memory consolidation across both age groups. Moreover, we observed that the identified scores of short- and long-delay brain profiles were lower in children in comparison to young adults. Thus, extending the conventional univariate analyses, our approach suggests that individual differences in short- and long-delay memory consolidation, which contain significant age-related variations, are associated with neural profiles comprised of distinct structural brain regions that are unique for short and long delays. In the following, we discuss each finding in detail.

### 4.1 Short- and long-delay memory consolidation

#### 4.1.1 Less robust short- and long-delay memory consolidation in children in comparison to young adults

Children showed steeper accuracy percentage change and thus lower short and long-delay RR in comparison to young adults, indicating reduced retained memory of prior-knowledge-dependent complex associative information across time. On the one hand, our result is not in line with the findings of higher short-delay memory consolidation (i.e., after one night of sleep) for incidental learning episodic tasks in 7-12-years-old children in comparison to young adults (Peiffer et al., 2020; Wang et al., 2018). These studies suggested that higher proportion of slow wave sleep in children in comparison to adults may contribute to possible age-related consolidation benefits. On the other hand, Wilhelm et al (2008) did not find a beneficial effect of sleep in 6-8-years-old children in comparison to young adults, showing a comparable consolidation performance after one night of sleep for declarative visuo-spatial and word-pairs associations. The mixed results may be attributed to the nature of memory measures. Namely, Wang et al. (2018) employed item memory and Peiffer et al. (2020) employed associative memory measures with stimuli that were not related to any prior knowledge. This might have precluded the possibility of semantic elaboration in conceiving prior-knowledge-based associations, as was utilized in our study and Wilhelm et al. (2008). Activated prior knowledge boosts memory consolidation for associative representations (Fernández & Morris, 2018; Tse et al., 2011). However, when pre-existing knowledge representations are less extensive or limited in children, it may counteract beneficial effects of sleep in children (Gaudreau et al., 2001; Ohayon et al., 2004) on the consolidation of newly correctly learnt associations. Therefore, lower short term retention rates in children, compared to adults, after one night of sleep in our study may be attributed to their lower level of prior knowledge for the stimuli. In similar fashion, superior long-delay memory consolidation in young adults may also be attributed to their more extensive world knowledge and, therefore, more elaborate schemas, ensuring better accessible memory representations in long term (Craik & Lockhart, 1972; Shing et al., 2010; van Kesteren et al., 2012).

On a related note, an adaptive, strategic learning of object-location associations that resulted in high level of final performance through repeated testing and encoding was applied in this study. It is worth noting that no pure measurement for consolidation was conducted, as this process happens offline after Day 0 learning. Essentially the consequences of consolidation at Day 1 and Day 14 retrieval were assessed. In view of this limitation, we opted for a 3-alternative forced choice procedure during retrieval to reduce the demand on retrieval processes (e.g., the need for strategic search in free recall). Adding to this, the retrieval procedure was kept comparable and stable over time, making sure that the process of retrieval was well-trained for all participants. Therefore, while we cannot entirely rule out variations in retrieval, by keeping the procedure the same across time we believe that our behavioral outcome largely revealed differences in consolidation process. In this way, in comparison to previous studies that aimed at episodic memory retrieval after one-shot encoding without mnemonic strategy use, our work shed some light on memory consolidation of well-studied information. Considering age differences in the final learning performance, we concentrated our analysis only on the consolidation of correctly learnt association. Our findings indicate that the potential beneficial effect of sleep for children (as reported in some studies) in comparison to adults for incidentally learned information may not apply to elaboratively learned information. This is potentially because such information is easier to be strengthened and integrated into the more extensive, well-connected network of knowledge in adults through sleep. It could also be remembered more easily by adults through more efficient strategic control of memories (e.g., self-generated cues) upon retrieval (Fandakova et al., 2017; Shing et al., 2008). In other words, the advantage of deliberate, repeated intentional learning, in comparison to accidental episodic learning, is more pronounced in adults in comparison to children (McDaniel & Masson, 1977), relying on the ability to utilize adequate learning operations (Eagle & Leiter, 1964). Taken together, our findings provide novel empirical evidence showing that, in the case of intentional encoding, 5-to-7-year-old children show less robust short- and long-delay memory consolidation of correctly learned object-location associations compared to young adults.

#### 4.1.2 Short- and long-delay memory consolidation are related to differential profiles of structural brain measures across both age groups

Based on the memory consolidation literature that postulates differential time-related neural reorganization of memory traces depending on the nature of stimuli (Moscovitch & Gilboa, 2021; Sekeres et al., 2017) and of learning (Gilboa & Marlatte, 2017), we expected to identify differential brain profiles that would reflect detail-rich memories in short delay and perceptually decayed mnemonic representations in long delay, which increase the relative importance of strategic control/elaboration for memory. Somewhat in line with our expectations, we identified a stable multivariate profile of short-delay memory consolidation comprised of neocortical (i.e., prefrontal, parietal, and occipital), cerebellar, and MTL (specifically, hippocampal subfields and EC) structures across age groups. The identified brain profile related to variations in long-delay memory consolidation is with a reduced number of brain regions, comprising of mostly neocortical regions of prefrontal, parietal and occipital cortex, as well as two specific hippocampal subfields, namely the subiculum and CA1-2.

First, our results extend previous univariate findings on the relations between brain measures and mnemonic processes in developmental cohorts. For example, studies showed that extended developmental trajectories of hippocampal subfields and EC (Canada et al., 2019; Keresztes et al., 2017), cerebellar (Sussman et al., 2016), prefrontal (Bauer et al., 2019; Botdorf & Riggins, 2018; Mills et al., 2016, 2016; Schlichting & Preston, 2015; Sousa et al., 2018; Sowell et al., 2001), parietal and occipital (DeMaster & Ghetti, 2013; Hebscher et al., 2019; Himmer et al., 2019; Karanian & Slotnick, 2015) regions are related to age-related differences in encoding and retrieval of memories. The brain profiles identified in our study extend the literature, showing that multivariate profiles comprised of these structural brain measures can also be related to memory retention across short and long delays. In particular, thinner medial OFC, IFG, rmPFC, LOC, PPC regions and larger volumes of hippocampal subfields, cerebellum und EC are associated with better memory consolidation after one night of sleep. The long delay brain profile shows that thinner IFG, OFC, rmPFC, PPC, and LOC as well as larger volumes of subiculum and CA1-2 hippocampal subfields are associated with better memory consolidation over two weeks. Notably, the directionality in these regions is in line with existing findings on developmental trajectories of brain morphology, i.e. thicker cortex and smaller MTL/cerebellum in 6-7-year-old children compared to adults (see Hedman et al., 2012 for an overview). This corresponds to the age-related and expected volumetric increase in cerebellum and hippocampus on the one hand, and cortical thinning in neocortical areas on the other hand. The derived brain scores also showed significant age difference, paralleling age-related differences in short- and long-delay RR. Taken together, our interpretation is that the brain profiles identified with PLSC may partly underlie children’s worse short- and long-delay consolidation compared to adults.

The distinctiveness of short- and long-delay brain profiles may be attributed as expected to time-related decay of detail-rich mnemonic representation. As our task required utilization of mnemonic strategies using prior knowledge to form vivid memories of object-location associations, we expected that the stabilization of memory traces for correctly learnt associations would depend on strategic elaborations based on prior knowledge and controlled processing. This should be the case both after one night of sleep as well as over longer time. On the other hand, detail-rich and strong mnemonic representation may be more prominent for short-delay than long-delay consolidation. The involvement of EC, hippocampal subfields, LOC and cerebellum in the short-delay profile is in agreement with our hypothesis, as these brain structures are important for perceptual vividness and precision of memory representations (DeMaster & Ghetti, 2013;

Fandakova et al., 2019; Grill-Spector et al., 2001; Karanian & Slotnick, 2015; Keresztes et al., 2017). There tends to be a decay of memory precision over longer time, which may explain why such regions as EC and DG-CA3 and cerebellum are no longer associated with variation in RR after two weeks. This is in line with Østby et al. (2012), who showed that variation in hippocampal volume was related to memory RR after one week and Fjell et al. (2019), who showed that memory RR over extended period of around 10 days was related to hippocampal and lateral prefrontal cortex structure. We did not expected, however, any associations of more fine-grained intrahippocampal structures with RR after more extended consolidation time of two weeks and due to associative nature of our task and lack of any developmental finding with this regard (Moscovitch & Gilboa, 2021). However, our finding of subiculum and CA1-2 hippocampal subfields being associated with age-related variations in long delay RR converges with recent evidence that in mice CA1 is necessary for long-delay consolidation of very remote memories or retrieval of gist memory, while CA3 is required for retrieval of precise memories recent in time (Atucha et al., 2021). In addition, Barry et al. (2021) showed a positive relationship between pre/parasubiculum volume and autobiographical memory over time, showing its role in the robustness of remote memory over time. In relation to this subiculum (Keresztes et al., 2022), and CA1 subfields (Riggins et al., 2018) was recently shown to undergo profound volumetric increase in middle childhood, indicating that the age-related increase in structural volume in these regions go hand in hand with improvement in long-delay memory consolidation.

Prefrontal (lateral and medial PFC), parietal and occipital brain structures, on the other hand, were associated with both short- and long-delay memory consolidation. With the decay of memory precision, the relative importance of these regions became even stronger. MOFC and rmPFC are associated with schema-integration (Brod & Shing, 2018; Mella et al., 2021; van Kesteren et al., 2012), while IFG and lOFC are associated with strategic elaboration and control over memories (Badre et al., 2005; Fjell et al., 2019; Kuhl et al., 2012; Østby et al., 2012). Presumably individuals with better profile in these regions could form memory representations in a controlled way, particularly by using prior knowledge for elaboration, leading to better memory performance both in short and long delay. This is in line with findings that age-related decrease in PFC volumes is related to increasing strategy use in cross-sectional sample of 5-25 year old participants (Yu et al., 2018). Furthermore, parietal regions such as PPC is found to be important for reinstating neural representation of visuo-spatial association (Brodt et al., 2016; DeMaster & Ghetti, 2013; Himmer et al., 2019; Takashima et al., 2007). In line with this, successful recollection of items with precise contextual details is found to be related to PPC in childhood cohort (DeMaster & Ghetti, 2013). Also LOC as constituent of both short and long delay brain profile is in line with its association with reinstatement of item-specific information upon retrieval (Grill-Spector et al., 2001; Karanian & Slotnick, 2015) and neural specificity of scene representation at retrieval (Fandakova et al., 2019), as out task despite decay of precision required associative location memory for both delays. Taken together, age-related differences in neocortical parietal and prefrontal brain regions, which are important for creating and accessing elaborative memory traces that are long lasting, may underlie children’s steeper decline in memory retention.

Finally, contrary to our expectation, PHC was not associated with memory RR at all. Despite PHG’s involvement in spatial context-related associative recollection (Ghetti et al., 2010; Ranganath & Ritchey, 2012), it is not involved in variations in short-delay memory consolidation of detail-rich visuo-spatial associations. Similarly, our findings show that, despite cerebellar involvement in declarative memory processes (Vecchi & Gatti, 2020), associative semantic memory for words (Gatti et al., 2021) and retrieval of long-term episodic memory (Andreasen et al., 1999), its structural volume is not linked to variations in long-delay memory consolidation within long-delay brain profile.

## 5 LIMITATIONS

Several limitations of the current study should be noted. First, despite our procedure of learning to the criterion to maximize comparability of retention rates, we observed group differences in initial memory performance. Future studies may incorporate individualized item-based approach of learning to criteria, excluding correctly remembered items from further learning cycles to ensure faster learning and lessening the overall task workload (Karpicke & Roedigeriii, 2007; McDermott & Zerr, 2019; Zerr et al., 2018). Alternatively, the task workload could be increased for the adults to ensure similar initial final learning performance. Second, we did not find reliable brain profile that relate to memory retention within the children and adult groups, respectively. This may be due to the narrow age range and restricted variation within each group, as our main questions call for maximizing between-group differences. Future studies may either extend the age range or increase sample sizes to create subgroups of high- and low-performers, allowing a clustering approach to look at differentiated neural profiles of variations in short- and long-delay memory consolidation. Third, the current findings concentrate mainly on associative memory of schema-congruent information. To investigate the beneficial effect of prior knowledge in memory consolidation, future studies should investigate how violations of knowledge, namely schema-incongruent information, may impact the learning of associative information and their subsequent consolidation in short- and long-delay memory and how this effect may differ throughout development. Fourth, we do not report sleep-related measures that may have an impact on memory consolidation. Finally, our sample of children was positively biased in IQ and mother’s education, in comparison to young adults. The former may just be because different IQ tests were used for children (WAIS-IV) and adults (K-ABC). The latter reflects generational difference in level of education. Nevertheless, the difference between children and adults in these aspects should be noted, as they may limit the generalizability of our results.

## 6 CONCLUSIONS

In this study, we provided novel empirical evidence that 5-to-7-year-old children consolidate intentionally learned object-location associations worse in comparison to young adults after one night of sleep and over an extended period of two weeks. We could identify distinct stable multivariate profiles comprised of specific memory-related brain regions that explain variation in either short- or long-delay memory consolidation. The brain regions involved support the notion that perceptually rich, vivid memory traces are important for variations in short delay, while controlled and elaboration processing are important for variations in both types of delay. As the memory consolidation shows strong relation to age and the identified brain profiles showed significant age differences, together the findings indicate that age-related differences in memory consolidation may be associated with specific maturational processes of distinct anatomically interconnected brain regions.

## Data availability statement

The datasets generated and analysed during the current study are available from the open science framework OSF under the following link: https://osf.io/qx2wp/.

## Acknowledgements

We thank all the children and parents who participated in the study. This work was supported by the Deutsche Forschungsgemeinschaft (DFG; German Research Foundation, Project-ID 327654276, SFB 1315, “Mechanisms and Disturbances in Memory Consolidation: from Synapses to Systems”). The work of YLS was also supported by the European Union (ERC-2018-StG-PIVOTAL-758898) and the Hessian Ministry of Higher Education, Science, Research and Art (Excellence Program, project ‘The Adaptive Mind’). The work of AMK was also supported by the Einstein Stiftung.

## Author contributions

Y.L.S, C.B., A.M.K secured funding. I.S and Y.L.S, C.B., A.M.K contributed to conception and design of the study. I.S., H.S., N.W.-C., and P.L. performed data collection and data curation. I.S., P.L., and F.P performed the statistical analysis. IS wrote the first draft of the manuscript, P.L., M.B. and F.P. wrote sections of the manuscript. All authors contributed to manuscript revision, read, and approved the submitted version.

