## Supplementary Materials for "Distinct Multivariate Structural Brain Profiles Are Related to Variations in Short- and Long-Delay Memory Consolidation Across Children and Young Adults"

### S.1 Supplementary Methods

#### S1.1. Stimuli selection and randomization

The semantic themes were grouped based on six overarching topics (e.g., nature, school, home, etc). Then four sets with 60 unique object-location associations each were created. In each set 30 semantic themes belonging to all six overarching topics were represented with two unique object-location associations. In this way, each set consisted of a balanced combination of semantic themes. One of these sets was randomly assigned to each participant. For each scene, we identified one out of the six possible previously defined placement areas as plausible for positioning the object into the scene. Within each placement area, objects could be placed in one of three possible locations, resulting in a total of 18 possible locations.

We applied a semi-random distribution of the stimuli from the batches between the two retrieval days. Out of 60 topics, 30 topics were learnt on Day 0, each topic presented by 2 different object-location pairs. Within one batch of stimuli, 30 object-scene pairs representing 15 different topics (2 pairs per topic) were presented in a random order. Moreover, four different version of such batches were created. Depending on the version of the task, two subsequent combinations for retrieval were possible. Irrespective of the random presentation order of pairs during learning, the first haft of the items from the 1st batch and the second half of the items from the 2nd batch were combined to create the 1 retrieval set. Then the 2nd haft of the 1st batch and the 1st half of the 2nd batch were combined to create the 2nd retrieval set. Depending on the version number and subversion number, these sets were semi-randomly appointed either for short-delay or long-delay retrieval and this appointment randomized across the participants.

S1.2. Assessment of demographic and cognitive covariates

Other cognitive covariate tasks were also assessed, such as cognitive switching and object-location memory with immediate test but were not included in the current paper.

***Day 0*:** After the experimental task, several subtests of the K-ABC II Test *(e.g., Atlantis, Rover, Rebus, Riddle and Atlantis delayed) were administered to children, while young adults were tested with the WAIS-IV Test.

***Day 1:*** In addition, children performed several subtests of the K-ABC II Test *(e.g., Expressive Vocabulary, Triangles, Pattern Reasoning). Day 14 In addition, children performed several subtests of the K-ABC II Test *(e.g., Patterns, Verbal Knowledge, Word Order).

#### S1.3. Structural Data Processing with Freesurfer Pipeline

The brain mask estimated previously was refined with a custom variation of the method to reconcile ANTs-derived and FreeSurfer-derived segmentations of the cortical gray-matter of Mindboggle (Klein et al., 2017). All data sets were transformed and organized according to the BIDS standard (Gorgolewski et al., 2016) using BIDSify (<https://github.com/MartinBa9210/BIDSify>). One T1-weighted (T1w) images was used within the input BIDS datasets. The T1-weighted (T1w) image was corrected for intensity non-uniformity (INU) with N4BiasFieldCorrection (Tustison et al., 2010), distributed with ANTs 2.3.3 (Avants et al., 2008; RRID:SCR_004757), and used as T1w-reference throughout the workflow. The T1w-reference was then skull-stripped with a Nipype implementation of the antsBrainExtraction.sh workflow (from ANTs), using OASIS30ANTs as target template. The brain mask estimated previously was refined with a custom variation of the method to reconcile ANTs-derived and FreeSurfer-derived segmentations of the cortical gray-matter of Mindboggle (RRID:SCR_002438, Klein et al., 2017). Volume-based spatial normalization to two standard spaces (MNI152NLin2009cAsym, MNI152NLin6Asym) was performed through nonlinear registration with antsRegistration (ANTs 2.3.3), using brain-extracted versions of both T1w reference and the T1w template. The following templates were selected for spatial normalization: ICBM 152 Nonlinear Asymmetrical template version 2009c (Fonov et al., 2009; RRID:SCR_008796; TemplateFlow ID: MNI152NLin2009cAsym), FSL’s MNI ICBM 152 non-linear 6th Generation Asymmetric Average Brain Stereotaxic Registration Model (Evans et al., 2012; RRID:SCR_002823; TemplateFlow ID: MNI152NLin6Asym).

#### S1.4. Structural Data Processing with ABCD Pipeline

PreFreesurfer normalizes anatomical data which entails brain extraction, denoising, and bias field correction on anatomical T1 weighted data. The ABCD-HCP pipeline includes two additional modifications to improve output image quality. ANTs (B. B. Avants et al., 2011) DenoiseImage models scanner noise as a Rician distribution and attempts to remove such noise from the T1 and T2 anatomical images. Additionally, ANTs N4BiasFieldCorrection attempts to smooth relative image histograms in different parts of the brain and improves bias field correction. (2) FreeSurfer (FreeSurfer 5.3.0; Dale et al., 1999) constructs cortical surfaces from the normalized anatomical data. This stage performs anatomical segmentation, white–grey and grey–CSF cortical surface construction, and surface registration to a standard surface template. Surfaces are refined using the T2 weighted anatomical data. Mid-thickness surfaces, which represent the average of white–grey and grey–CSF surfaces, are generated. (3) PostFreesurfer converts prior outputs into an HCP-compatible format (that is, CIFTIs) and transforms the volumes to a standard volume template space using ANTs nonlinear registration, and the surfaces to the standard surface space via spherical registration.(B. B. Avants et al., 2011; Dale et al., 1999).

### S2. Supplementary Results

#### S2.1. Results based on all correct items learned on Day 0.

The memory retention rates for Day 1 and Day 14 used for the following analysis are relative to all correct items learnt on Day 0. Final learning performance is identical for everyone and used as a baseline for the memory consolidation.

**Table S1.** *Statistical overview of the main and interaction effects of the linear mixed effects model with subjects as random intercepts and the fixed effects of Group (children and young adults), Session (Day 1, Day 14, relative to correctly recalled items on Day 0), IQ, Handedness, and Group x Session interaction on the memory retention rates.*

|  | *actrial* | | |
| --- | --- | --- | --- |
| *Predictors* | *Estimates* | *CI* | *p* |
| *(Intercept)^a^* | 100.60 | 96.10 – 105.10 | **<.001** |
| Short Delay (Session, Day1) | -16.63 | -19.77 – -13.49 | **<.001** |
| Long Delay (Session, Day14) | -38.57 | -41.71 – -35.43 | **<.001** |
| Group | .52 | -3.19 – 4.24 | .783 |
| IQ | .10 | .01 – .19 | **.033** |
| Sex | 2.58 | -.17 – 4.98 | **.036** |
| Handedness | -1.84 | -5.29 – 1.60 | .295 |
| Group * Short Delay | 11.25 | 6.51 – 15.99 | **<.001** |
| Group * Long Delay | 14.73 | 9.99 – 19.47 | **<.001** |
| **Random Effects** | | | |
| σ2 | 64.17 | | |
| τ00 subject | 10.44 | | |
| ICC | 0.14 | | |
| N subject | 89 | | |
| Observations | 267 | | |
| Marginal R2 / Conditional R2 | 0.737 / 0.774 | | |

*Notes.*^a^The following reference levels where used: for session, Day0; for Group, Children; for Sex, male; for Handedness, right hand. IQ = Intelligence Quotient based K-ABC (Kaufman & Kaufman, 2012) for children and WAIS-IV (Wechler, 2012) for young adults; σ2 – residuals, τ00 – variance of the random intercept. Type III Analysis of Variance Table with Satterthwaite's method. *p < .05; ** <.01, ***<.001 (significant difference).

#### S2.2. Results based on all correct items learned on Day 0 for participants who reached set learning criteria of 83% during final learning.

The memory retention rates for Day 1 and Day 14 used for the following analysis are relative to all correct items learnt on Day 0 for participants who reached set learning criteria of 83% during final learning. Final learning performance is identical for everyone and used as a baseline for the memory consolidation.

**Table S2.** *Descriptive statistics of memory performance based on percentage of correct answers by age groups for participants who fulfilled the learning criteria of 83% correct responses.*

| **Session** | Children  (n = 42) | | Young Adults  (n = 39) | | Group effect  (CH vs YA) | |
| --- | --- | --- | --- | --- | --- | --- |
|  | *M* | *SD* | *M* | *SD* | *p-value* | *w^2^* |
| **Retrieval** |  |  |  |  |  |  |
| Day 0 ^average^ | 1 | 0 | 1 | 0 | - | - |
| Day 1* | 83.94 | 12.02 | 94.62 | 5.90 | *** | .24 |
| Day 14* | 60.98 | 11.98 | 76.16 | 12.03 | *** | .28 |

*Notes.** Relative to all correct items on Day 0 for all participants who reached the initially set criteria of 83 % correct responses in final learning. M = mean; SD = standard deviation.

**Table S3.** *Statistical overview of the main and interaction effects of the linear mixed effects model with subjects as random intercept and the fixed effects Group (children and young adults), Session (Day 1, Day 14, relative to correctly recalled items on Day 0 for participants who reached set learning criteria of 83% during final learning), IQ, Handedness, and Group x Session interaction on the memory retention rates.*

|  | *actrial* | | |
| --- | --- | --- | --- |
| *Predictors* | *Estimates* | *CI* | *p* |
| *(Intercept)^a^* | 100.42 | 95.68 – 105.16 | **<.001** |
| Short Delay (Session, Day1) | -16.06 | -19.38 – -12.74 | **<.001** |
| Long Delay (Session, Day14) | -39.02 | -42.34 – -35.70 | **<.001** |
| Group | .69 | -3.15 – 4.54 | .724 |
| IQ | .10 | .00 – .20 | **.047** |
| Sex | 2.49 | -.03 – 5.01 | **.053** |
| Handedness | -1.79 | -5.29 – 1.71 | .32 |
| Group * Short Delay | 10.68 | 5.90 – 15.46 | **<.001** |
| Group * Long Delay | 15.18 | 10.40 – 19.96 | **<.001** |
| **Random Effects** | | | |
| σ2 | 60.16 | | |
| τ00 subNo | 12.46 | | |
| ICC | 0.17 | | |
| N subNo | 81 | | |
| Observations | 243 | | |
| Marginal R2 / Conditional R2 | 0.740 / 0.785 | | |

*Notes.* ^a^The following reference levels where used: for session, Day0; for Group, Children; for Sex, male; for Handedness, right hand. IQ = Intelligence Quotient based K-ABC (Kaufman & Kaufman, 2012) for children and WAIS-IV (Wechler, 2012) for young adults; σ2 – residuals, τ00 – variance of the random intercept. Type III Analysis of Variance Table with Satterthwaite's method. *p < .05; ** <.01, ***<.001 (significant difference).

#### S2.3. Results based on all items learned on Day 0.

The following results are relative to all learnt items on Day 0.

**Table S4.** *Descriptive statistics of memory performance based on percentage of correct answers by age groups.*

| **Session** | Children  (n ­= 50) | | Young Adults  (n = 39) | | Group effect  (CH vs YA) | |
| --- | --- | --- | --- | --- | --- | --- |
|  | *M* | *SD* | *M* | *SD* | *p-value* | *w^2^* |
| **Retrieval** |  |  |  |  |  |  |
| Day 0* | .908 | .043 | .979 | .027 | **** | .48 |
| Day 1 | .796 | .129 | .939 | .067 | **** | .30 |
| Day 14 | .594 | .121 | .760 | .119 | **** | .32 |

*Notes.* * Averaged across all trials learnt on Day 0. M = mean; SD = standard deviation.

**Table S5.** *Statistical overview of the main and interaction effects of the linear mixed effects model with subjects as random intercept and the fixed effects Group (children and young adults), Session (Day 0, Day 1, Day 14), IQ, Handedness, and Group x Session interaction on the memory retention rates.*

| *Predictors* | *Estimates* | *95% CI* | *p-value* |
| --- | --- | --- | --- |
| (Intercept)^a^ | .91 | .86 – .96 | **< .001** |
| Short Delay (Session, Day 1) | -.11 | -.14 – -.08 | **< .001** |
| Long Delay (Session, Day 14) | -.31 | -.35 – -.28 | **< .001** |
| Group | .08 | .04 – .12 | **< .001** |
| IQ | .00 | .00 – .00 | **.034** |
| Sex | .03 | .00 – .06 | **.049** |
| Handedness | -.01 | -.05 – .02 | .451 |
| Group * Short Delay | .07 | .03 – .12 | .**003** |
| Group * Long Delay | .09 | .05 – .14 | **< .001** |
| **Random Effect** | | | |
| σ^2^ | .01 | | |
| τ_00_ _subNo_ | .00 | | |
| ICC | .25 | | |
| N _subNo_ | 89 | | |
| Observations | 267 | | |
| Marginal R^2^ / Conditional R^2^ | 0.684 / 0.762 | | |

*Notes.* ^a^The following reference levels where used: for session, Day0; for Group, Children; for Sex, male; for Handedness, right hand. IQ = Intelligence Quotient based K-ABC (Kaufman & Kaufman, 2012) for children and WAIS-IV (Wechler, 2012) for young adults; σ2 – residuals, τ00 – variance of the random intercept. Type III Analysis of Variance Table with Satterthwaite's method. *p < .05; ** <.01, ***<.001 (significant difference).

#### S2.4. Results based on matched on Day 0 performance of adult-child dyads separately for short- and long-delay.

**Table S6.** *Statistical overview of the main and interaction effects of the linear mixed effects model with subjects as random intercept and the fixed effects Group (children and young adults), Session (Day 1 relative to correct items on Day 0), IQ, Handedness, and Group x Session interaction on the memory retention rates.*

| *Predictors* | *Estimates* | *95% CI* | *p-value* |
| --- | --- | --- | --- |
| (Intercept)^a^ | 99.98 | 94.98 – 104.97 | **< .001** |
| Short Delay (Session, Day 1) | -16.51 | -20.32 – -12.71 | **< .001** |
| Group | -.40 | -4.29 – 3.49 | **< .001** |
| IQ | .05 | -.05 – .16 | .335 |
| Sex | 2.95 | .15 – .5.76 | **.039** |
| Handedness | -.89 | -4.80 – 3.02 | .655 |
| Group * Short Delay | 9.89 | 4.51 – 15.28 | <.**001** |
| **Random Effect** | | | |
| σ^2^ | 43.38 | | |
| τ_00_ _subNo_ | .00 | | |
| N _subNo_ | 46 | | |
| Observations | 92 | | |
| Marginal R^2^ | 0.529 | | |

*Notes.* ^a^The following reference levels where used: for session, Day0; for Group, Children; for Sex, male; for Handedness, right hand. Adult and children’s participants were matched based on their final learning performance on Day 0. It resulted in 28 matched child-adult dyads for short delay retrieval. IQ = Intelligence Quotient based K-ABC (Kaufman & Kaufman, 2012) for children and WAIS-IV (Wechler, 2012) for young adults; σ2 – residuals, τ00 – variance of the random intercept. Type III Analysis of Variance Table with Satterthwaite's method. *p < .05; ** <.01, ***<.001 (significant difference).

**Table S7.** *Statistical overview of the main and interaction effects of the linear mixed effects model with subjects as random intercept and the fixed effects Group (children and young adults), Session (Day 14 relative to correct items on Day 0), IQ, Handedness, and Group x Session interaction on the memory retention rates.*

| *Predictors* | *Estimates* | *95% CI* | *p-value* |
| --- | --- | --- | --- |
| (Intercept)^a^ | 97.25 | 90.26 – 104.25 | **< .001** |
| Long Delay (Session, Day 14) | -37.89 | -43.42 – -32.36 | **< .001** |
| Group | .86 | -5.35 – 7.07 | .786 |
| IQ | .04 | -.14 – .21 | .690 |
| Sex | 2.71 | -1.41 – 6.83 | .198 |
| Handedness | .76 | -3.75 – 5.27 | .743 |
| Group * Long Delay | 8.58 | .76 – 16.40 | **.031** |
| **Random Effect** | | | |
| σ^2^ | 59.66 | | |
| τ_00_ _subNo_ | .00 | | |
| N _subNo_ | 30 | | |
| Observations | 60 | | |
| Marginal R^2^ | 0.833 | | |

*Notes.* ^a^The following reference levels where used: for session, Day0; for Group, Children; for Sex, male; for Handedness, right hand*.* Adult and children’s participants were matched based on their final learning performance on Day 0. It resulted in 15 matched child-adult dyads for short delay retrieval. IQ = Intelligence Quotient based K-ABC (Kaufman & Kaufman, 2012) for children and WAIS-IV (Wechler, 2012) for young adults; σ2 – residuals, τ00 – variance of the random intercept. Type III Analysis of Variance Table with Satterthwaite's method. *p < .05; ** <.01, ***<.001 (significant difference).

#### S2.5. Results based on structural brain measures.

**Table S8.** *Results of the analyses on group differences for brain structural measures.*

| *Region* | *Volume* | *Thickness* | *Children* | *Young Adults* | *t_(df)_* | *95% CI* | *p-value* | |
| --- | --- | --- | --- | --- | --- | --- | --- | --- |
|  |  |  | *Mean* | |  |  |  |  |
| Entorhinal cortex | *x* |  | *584.7* | *668.7* | *-4.24_(75,71)_* | *-123.54 – -44.55* | *<.0001* | ****** |
| Subiculum | *x* |  | *673.9* | *818.4* | *-7.43_(63,14)_* | *-183.28 – -105.61* | *<.0001* | ****** |
| *Dental Gyrus* | *x* |  | *481.7* | *589.9* | *-5.5747_(60.21)_* | *-146.96 – -69.35* | *<.0001* | ****** |
| *CA1-2* | *x* |  | *384.1* | *478.3* | *-7.3828_(71.50)_* | *-119.64 – -68.76* | *<.0001* | ****** |
| *Cerebellum*† | *x* |  | *110915.4* | *119966.5* |  | *4286 – 13816* | *0.000359* | ***** |
| *Parsopercularis* |  | *x* | *6.136* | *5.451* | *11.209_(73.52)_* | *0.56– 0.81* | *<.0001* | ****** |
| *Parsorbitalis* |  | *x* | *6.656* | *5.717* | *10.996_(76.58)_* | *0.77 – 1.11* | *<.0001* | ****** |
| *Parstriangularis*† |  | *x* | *5.994* | *5.196* |  | *-0.92 – -0.67* | *<.0001* | ****** |
| *Rostralmiddlefrontal*† |  | *x* | *5.719* | *4.796* |  | *-1.03 – -0.81* | *<.0001* | ****** |
| *Lateralorbitofrontal* |  | *x* | *6.467* | *5.652* | *13.773_(78.30)_* | *0.70 – 0.93* | *<.0001* | ****** |
| *Superiorparietal*† |  | *x* | *5.202* | *4.615* |  | *-0.70 – -0.48* | *<.0001* | ****** |
| *Medialorbitofrontal* |  | *x* | *6.101* | *4.962* | *17.255_(69.15)_* | *1.01 – 1.27* | *<.0001* | ****** |
| *Inferiorparietal*† |  | *x* | *5.896* | *5.277* |  | *-0.74 – -0.50* | *<.0001* | ****** |
| *Precuneus*† |  | *x* | *5.853* | *5.065* |  | *-0.90 – -0.67* | *<.0001* | ****** |
| *Parahippocampal* |  | *x* | *6.508* | *6.096* | *3.497_(76.80)_* | *0.18 – 0.65* | *0.0007855* | ****** |
| *Lateraloccipital* |  | *x* | *4.975* | *4.613* | *6.5995_(78.95)_* | *0.25 – 0.47* | *<.0001* | ****** |

*Notes.* Statistical values are shown for the t-test*.* † In case of violated assumptions of homogeneity of variances, a Games-Howell test was performed (Lee & Lee, 2018). CI – confidence interval; t – t-value; df – degrees of freedom; p-value: *p < .05; ** <.01, ***<.001 (significant difference), uncorrected for multiple comparisons; ns: non-significant difference.

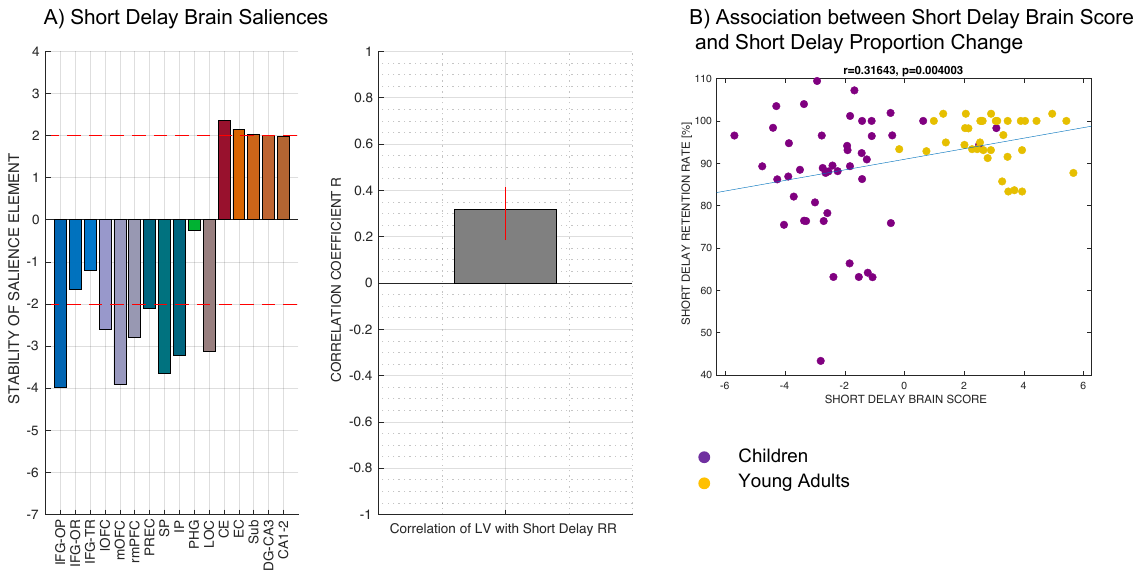

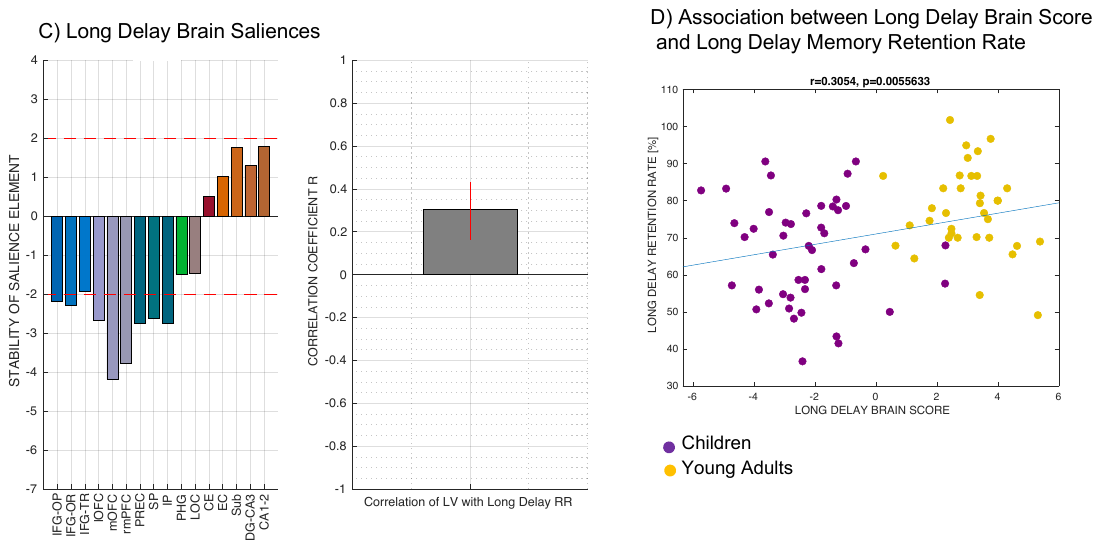

**Figure S1. Multivariate profiles of brain structures associated with variations in memory consolidation relative to all items learnt on Day 0.** A) *Short Delay Brain Saliences*. Brain saliences or latent variables weights for each ROI were incorporated in the analysis to create one latent variable that expresses a composite short-delay brain score. Stability of salience elements is defined by Z-scores (depicted as red line: a value larger/smaller than ±1.96 is treated as reliably robust at (a < .05). B) *Association between Short Delay Brain Score and Short Delay Retention Rate*. Brain short delay score or a latent variable plotted against short delay retention rate. Little overlap of brain short delay score between age groups (purple for children, yellow for young adults) shows that there are age-related differences in brain short delay scores.

C) *Long Delay Brain Saliences*. Brain saliences or latent variables weights for each ROI were incorporated in the analysis to create one latent variable that expresses a composite long-delay brain score. Stability of salience elements is defined by Z-scores (depicted as red line: a value larger/smaller than ±1.96 is treated as reliably robust at (a < .05). D) *Association between Long Delay Brain Score and Long Delay Retention Rate*. Brain long delay score or a latent variable plotted against short delay retention rate. Little overlap of brain long delay score between age groups (defined in purple (children) and yellow (young adults)) shows that there are age-related differences in brain long delay score.

*Note:* IFG – inferior frontal gyrus; OP – parsopercularis; OR – parsorbitalis; TR – parstriangularis; lOFC – lateral orbitofrontal cortex; mOFC – medial orbitofrontal cortex; rmPFC – rostromedial cortex; PREC – precuneus; SP – superior parietal cortex; IP – inferior parietal cortex; PHG – parahippocampal cortex; LOC – lateral occipital cortex; CE – cerebellum; EC – entorhinal cortex; SUB – Subiculum; DG-CA3 – dental gyrus and CA3; CA1-2 – CA1 and CA2 subfields of hippocampus. For group differences for each structural ROI (Houston et al., 2013), please refer to Table S8 in Supplementary Materials.
